# Identification of Differentially Expressed Genes in Wheat NILs in Response to Leaf Rust Infection using In Silico Analysis

**DOI:** 10.1101/2023.03.28.534481

**Authors:** Rahul Kumar

## Abstract

In this study, the molecular basis of leaf rust resistance was investigated by examining the differential gene expression of wheat cultivar Thatcher, both with and without a leaf rust resistance gene, in response to rust infection. unigenes/ESTs databases were utilized belonging to three near-isogenic lines (NILs) of Thatcher, including Thatcher, Thatcher + Lr10, and Thatcher + Lr1, to identify differentially expressed unigenes (DEUs) in response to leaf rust. Sixty-eight DEUs were identified using a data mining tool called Digital Differential Display (DDD). Among these DEUs were specific unigenes associated with Lr1 and/or Lr10-mediated resistance, as well as unigenes expressed exclusively in compatible interactions. Our results suggest that photosynthesis, lignin, and antifungal proteins play a role in the leaf rust resistance mechanism, as indicated by the overexpression of Ribulose-1,5-bisphosphate carboxylase/oxygenase, caffeic acid O-methyltransferase, and Glutathione S-transferase unigenes, respectively. Our findings demonstrate both common and distinctive genetic pathways involved in rust resistance in the Lr1 and Lr10 NILs. The expression of most DEUs was found to be high in leaves, moderate in sheath, stem, and inflorescence, and low in seed, root, flower, crown, callus, and cell culture, indicating their role in leaf rust. Using high-throughput microarray gene expression data, a subset of DEUs was validated, and as a result, eleven unigenes were mapped to deletion bins on individual wheat chromosomes. These key genes may be used as molecular markers for genome editing, which could facilitate the development of leaf rust-resistant wheat cultivars.

## Introduction

Leaf rust, caused by the fungal pathogen *Puccinia triticina*, is a significant threat to global wheat production, causing substantial economic losses (Dinh et al., 2020). Developing rust-resistant wheat cultivars is the most effective way to manage this disease. However, the development of such cultivars is a challenging task due to the complex and dynamic interactions between the host plant and the pathogen. Plant defense mechanisms against pathogens involve the expression of a wide range of genes, and identifying these differentially expressed genes can provide important insights into the underlying molecular mechanisms of plant defense against pathogens.

The recognition of pathogens by host plants is mediated by two classes of receptors, pattern recognition receptors (PRRs) and resistance (R) proteins, triggering two corresponding immune responses known as PAMP-triggered immunity (PTI) and effector-triggered immunity (ETI) (Jones and Dangl, 2006; Dodds and Rathjen, 2010). PTI is a basal defense response that occurs upon recognition of conserved microbial molecules known as pathogen-associated molecular patterns (PAMPs) by PRRs. This response results in the induction of various defense mechanisms, such as the production of reactive oxygen species (ROS), calcium ion influx, and changes in gene expression, which are aimed at halting pathogen growth and spread (Zipfel, 2014). However, some pathogens have evolved to overcome PTI through the secretion of effectors that can modify host cell processes, leading to the activation of ETI. ETI typically involves a rapid and robust defense response, characterized by the programmed cell death of infected host cells and the systemic activation of defense genes, leading to the establishment of long-lasting resistance (Jones and Dangl, 2006; Dodds and Rathjen, 2010). The R proteins are encoded by resistance genes (R genes), which are highly diverse and can confer resistance to a broad range of pathogens (Cui et al., 2015; Kourelis and van der Hoorn, 2018).

Pathogenesis-related (PR) proteins, such as chitinases and glucanases, are also known to play an important role in plant defense against fungal pathogens (Kumar et al., 2019; Singh et al., 2019; El-Kereamy et al., 2019). Chitinases are enzymes that degrade chitin, a component of fungal cell walls, while glucanases degrade β-1,3-glucans, another major component of fungal cell walls. Several studies have shown that the expression of chitinase and glucanase genes is induced in wheat in response to leaf rust infection (Liu et al., 2015; Singh et al., 2015). In addition, the overexpression of chitinase genes has been shown to enhance wheat resistance to leaf rust (Jha et al., 2016). Photosynthesis is also known to play an important role in plant defense against fungal pathogens. Several studies have shown that photosynthesis is inhibited in wheat in response to leaf rust infection (Liu et al., 2018; Yan et al., 2016). This inhibition of photosynthesis is thought to be due to the redirection of energy and resources to defense-related processes, such as the synthesis of defense-related proteins and phytohormones.

Hormones, such as salicylic acid (SA) and jasmonic acid (JA), are also known to play an important role in plant defense against fungal pathogens (Pieterse et al., 2012; Wang & Zhang, 2018; Vlot et al., 2009; Kazan and Lyons, 2014; Liu and Liu, 2021). SA is involved in the regulation of the defense response against biotrophic pathogens, while JA is involved in the regulation of the defense response against necrotrophic pathogens. Several transcriptome studies have shown that the expression of genes involved in the biosynthesis and signaling of SA and JA is induced in wheat in response to leaf rust infection (Chen et al., 2013; Liu et al., 2018). In addition, the exogenous application of SA and JA has been shown to enhance wheat resistance to leaf rust (Liu et al., 2018). Antifungal proteins, such as thionins, defensins, and lipid transfer proteins (LTPs), are also known to play an important role in plant defense against fungal pathogens. Several studies have shown that the expression of genes encoding for antifungal proteins is induced in wheat in response to leaf rust infection (Liu et al., 2018; Yan et al., 2016). In addition, the overexpression of antifungal protein genes has been shown to enhance wheat resistance to leaf rust (Jha et al., 2017).

Disease responses involve a coordinated differential expression of a large number of genes, which can be examined by any one of the three well known approaches, namely ‘analog’, ‘digital’, and ‘PCR-based’ approaches (Audic and Claverie 1997, Higuchi *et al*. 1992). Functional genomics approaches have provided a wealth of data on gene expression in response to various biotic and abiotic stresses in plants. These data sets, including microarray, MPSS, ESTs, and SAGE, can be used for in silico analysis of gene expression, allowing for identification of differentially expressed genes. Data mining tools such as Digital Differential Display (DDD), cDNA profiler, cDNA Digital Gene Expression Displayer (DGED), and SAGE map are available to facilitate identification of these differentially expressed genes (Strausberg et al. 2001). Digital Differential Display (DDD) is a method used to analyze gene expression and has been used extensively in plants (Yu et al., 2021; Cao et al., 2019; Chen et al., 2019; Gao et al., 2019; Huang et al., 2019). UniGene database is a valuable resource for in silico identification of differentially expressed genes, as it represents a non-redundant set of gene-oriented clusters derived from ESTs (Wheeler et al. 2004).

In this study, we used UniGene database to identify differentially expressed wheat UniGenes in response to leaf rust inoculation. These UniGenes were further validated using microarray gene expression data available at PLEXdb database. EST Profile database was used to quantify the expression of these UniGenes in different wheat tissues. Furthermore, a few validated UniGenes were physically mapped to individual wheat chromosomes using mapped ESTs/EST contigs available at GrainGenes 2.0 database. This approach provides a comprehensive analysis of gene expression in response to leaf rust in wheat and may help to identify key genes involved in leaf rust resistance.

## Materials and Methods

### Datasets used for digital gene expression analysis

For the identification of UniGenes that respond to leaf rust infection in wheat, two online datasets were utilized for digital gene expression analysis. These datasets, wheat UniGene and EST Profile, were obtained from NCBI. The wheat UniGene dataset consisted of three cDNA libraries named A (21046), B (21047), and C (8939) (Table 1). To obtain the EST Profile dataset for each differentially expressed UniGene (DEU), the corresponding web page of that particular UniGene in EST Profile Viewer was retrieved.

**Table 1.**
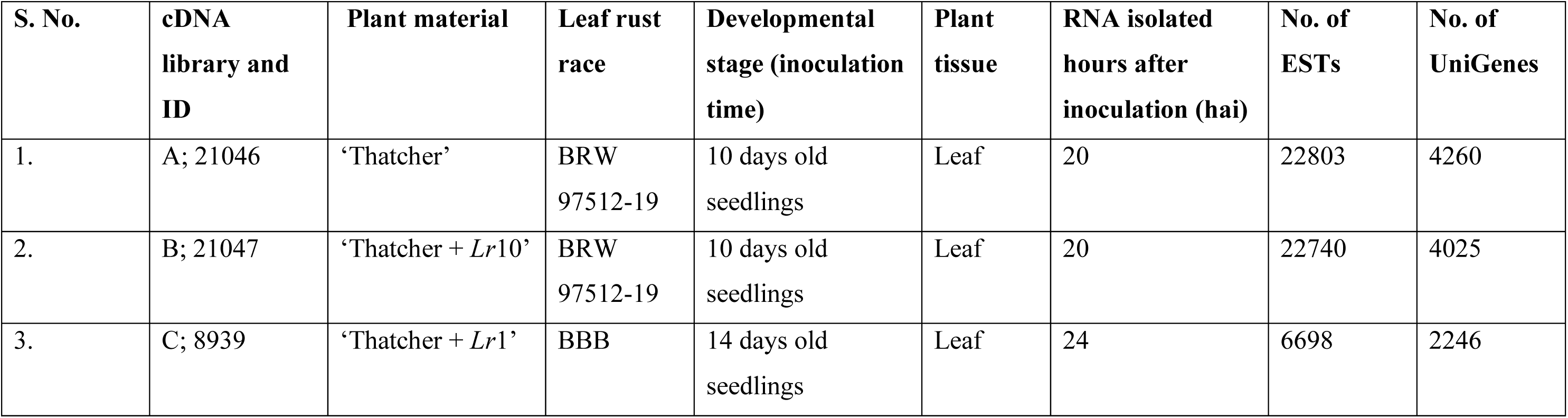
A summary of the details of UniGenes used for digital differential display, along with data sets of corresponding ESTs and cDNA libraries from which these ESTs were derived.

| **S. No.** | **cDNA library and ID** | **Plant material** | **Leaf rust race** | **Developmental stage (inoculation time)** | **Plant tissue** | **RNA isolated hours after inoculation (hai)** | **No. of ESTs** | **No. of UniGenes** |
| --- | --- | --- | --- | --- | --- | --- | --- | --- |
| 1. | A; 21046 | ‘Thatcher’ | BRW<br>97512-19 | 10 days old seedlings | Leaf | 20 | 22803 | 4260 |
| 2. | B; 21047 | ‘Thatcher + *Lr10*’ | BRW<br>97512-19 | 10 days old seedlings | Leaf | 20 | 22740 | 4025 |
| 3. | C; 8939 | ‘Thatcher + *Lr1*’ | BBB | 14 days old seedlings | Leaf | 24 | 6698 | 2246 |

### Wheat tissues for digital differential display analysis

The tissue-specific digital differential display analysis was conducted using ESTs from ten (10) different wheat tissues, namely callus, cell culture, crown, flower, inflorescence, leaf, root, leaf sheath, seed, and stem. The numbers of ESTs available for each of these tissues are provided in Table 2.

**Table 2.** Details of the number of ESTs present in each of the ten tissues used for Digital Differential Display analysis.

| **S. No.** | **Tissue** | **Number of ESTs** |
| --- | --- | --- |
| 1. | Callus | 10609 |
| 2. | Cell Culture | 10164 |
| 3. | Crown | 13685 |
| 4. | Flower | 64599 |
| 5. | Inflorescence | 121335 |
| 6. | Leaf | 57699 |
| 7. | Root | 167008 |
| 8. | Seed | 162266 |
| 9. | Sheath | 1102 |
| 10. | Stem | 93668 |

### Digital gene expression analysis

The digital gene expression analysis was performed using the Digital Differential Display (DDD) program available in the UniGene database of NCBI. The program utilizes the Fisher Exact test (Siegel 1956; Eujayl and Morris 2009) in combination with the Bonferroni inequality test to determine statistically significant differences in the abundance of a particular sequence between two different libraries. These tests are particularly suitable for analyzing randomly selected genes from pools or libraries of unequal size, and to decipher statistical significance.

To identify differentially expressed genes in the wheat host plant in response to pathogen inoculation, DDD was used to compare two members of each of the following three pairs of hosts: (i) Thatcher and Thatcher + Lr10, (ii) Thatcher and Thatcher + Lr1, and (iii) Thatcher + Lr10 and Thatcher + Lr1. Additionally, the expression of each differentially expressed UniGene was quantified in 10 different wheat tissues using the available EST Profile dataset. The actual numbers of ESTs belonging to each UniGene were represented as a fraction of all ESTs in a cDNA library, and the data were normalized by converting the fraction into transcripts per million (TPM) sequences. Fisher’s exact test was performed using the actual number of ESTs with a cutoff value of P ≤ 0.05, and a minimum of one EST of a UniGene in a treatment/library was considered as a threshold to understand the expression of a UniGene.

The EST frequency (in TPM) of an individual UniGene was calculated using the following formula: TPM = [Number of ESTs belonging to a UniGene/Total number of ESTs in the library] × 1,000,000. The fold difference in expression of an individual UniGene in two libraries (e.g. library A and B, A and C, B and C) was calculated using the following formula: Fold difference = [(TPM in library 1)/(TPM in library 2)].

### Cluster analysis

The expression data for clustering generation were based on the frequencies of ESTs belonging to individual UniGene, represented as TPM values. Hierarchical clustering of differentially expressed UniGenes (DEUs) was performed using complete linkage analysis, which calculates the distance between two clusters as the maximum distance between a pair of objects, one in one cluster and one in the other. Pearson correlation coefficients were used to calculate the distance. The analysis was conducted using Cluster 3.0 software downloaded from the University of Tokyo website.

To cluster different tissues and DEUs based on their expression profiles, average linkage analysis was performed. This method calculates the distance between two clusters as the average distance between objects from the first cluster and objects from the second cluster. Each pool was given equal weight to normalize the number of libraries in each pool. The results were visualized in graphical form using *.CDT, *.GTR, and *.ATR files generated by Cluster 3.0.

### Validation of expression pattern of UniGenes

To identify potential microarray probes that could represent the DEUs identified using DDD, a Blast search was performed using the nucleotide sequences of all DEUs against the probes of the Wheat 61K microarray chip available in PLEXdb (Plant Expression Database; http://www.plexdb.org/plex.php?database=Wheat). Sequences that showed an E-value of 0.0 and >95% similarity with the microarray probes were selected. The gene expression data of two experiments, TA3 (transcription patterns during wheat development) and TA32 (transcript profiling of Lr1- and Lr34-mediated leaf rust resistance in wheat), were retrieved for these selected probes. From experiment TA32, the expression data of wheat leaf rust-resistant gene Lr1 was downloaded and compared with the results obtained from DDD analysis.

### In-silico physical mapping of differentially expressed wheat UniGenes

In silico physical mapping was conducted to determine the chromosomal locations of UniGenes that showed significant matches with wheat microarray probes. The nucleotide sequences of these UniGenes were Blast searched against mapped ESTs/mapped EST contigs available at the GrainGenes 2.0 website. UniGenes that showed >95% similarity and had an E-value from 0.0 to 1E-10 were considered to have a significant match and were placed in the corresponding deletion bins on wheat chromosomes. The distribution of loci among the three subgenomes, the seven homoeologous groups, and chromosomes arms was tested using the χ2 test for goodness-of-fit. The expected number of loci was estimated based on the physical length (in micrometers) and DNA contents of wheat chromosomes and chromosome arms, assuming a random distribution of loci among the three subgenomes, seven homoeologous groups, and among chromosome arms.

## Results and Discussion

### Identification of DEUs

In this study, we used DDD to analyze the expression of genes in wheat NILs inoculated with leaf rust using wheat ESTs available in three cDNA libraries, each belonging to a different wheat NIL. We identified a total of 68 UniGenes showing differential expression across the three NILs. While 10 UniGenes were unique to an individual NIL, the number of DEUs did not differ significantly across the three NILs. The highest number of UniGenes showing differential expression was observed in Thatcher + Lr10, followed by Thatcher + Lr1 and Thatcher. However, the available ESTs in the three libraries differed significantly, indicating that the number of UniGenes differentially expressed in a NIL was not directly related to the total number of ESTs in the corresponding library, but instead varied with the type of treatment.

We determined the level of expression of each UniGene in a specific NIL by the number of ESTs representing that UniGene in the cDNA library belonging to an individual NIL. The total number of ESTs belonging to an individual UniGene that was differentially expressed ranged from 23 ESTs to 13610 ESTs, with an average of 768.25 ESTs per DEU (Table 3). The range of ESTs belonging to an individual UniGene varied from 1 to 1000 for two-thirds of the DEUs, suggesting that at least two-thirds of UniGenes belonged to the class in which the mean is situated, and the set of UniGenes used was suitable for the study of comparative profiling of ESTs (Table 3).

**Table 3.**
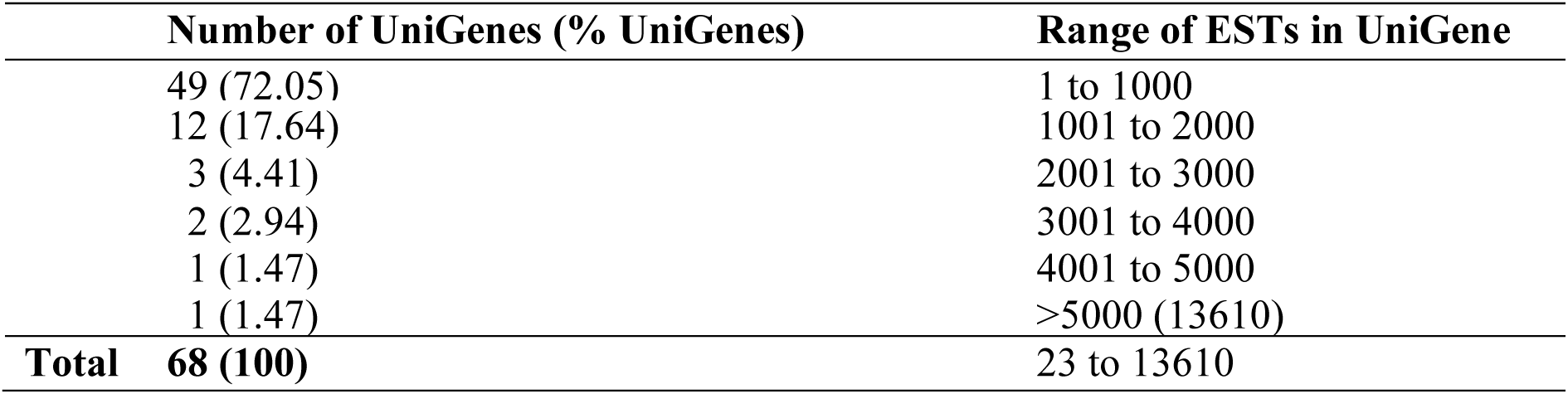
Number of differentially expressing UniGenes and the range of the number of ESTs in the corresponding UniGenes.

Out of the 68 differentially expressed UniGenes, 43 showed expression across all three NILs, although the difference in the level of their expression between any two NILs varied many folds. The remaining 25 UniGenes were expressed in only one (10 UniGenes) or two (15 UniGenes) NILs. Functional annotation of these UniGenes revealed that they were involved in housekeeping, abiotic stress tolerance, and disease resistance. We classified the 68 DEUs as follows: (i) genes showing high expression in the susceptible host Thatcher, (ii) genes showing high expression in the resistant host(s), and (iii) genes with non-specific expression. Table 4 provides details of UniGenes that expressed differentially across all three NILs, including their annotation and fold change in their expression.

**Table 4.**
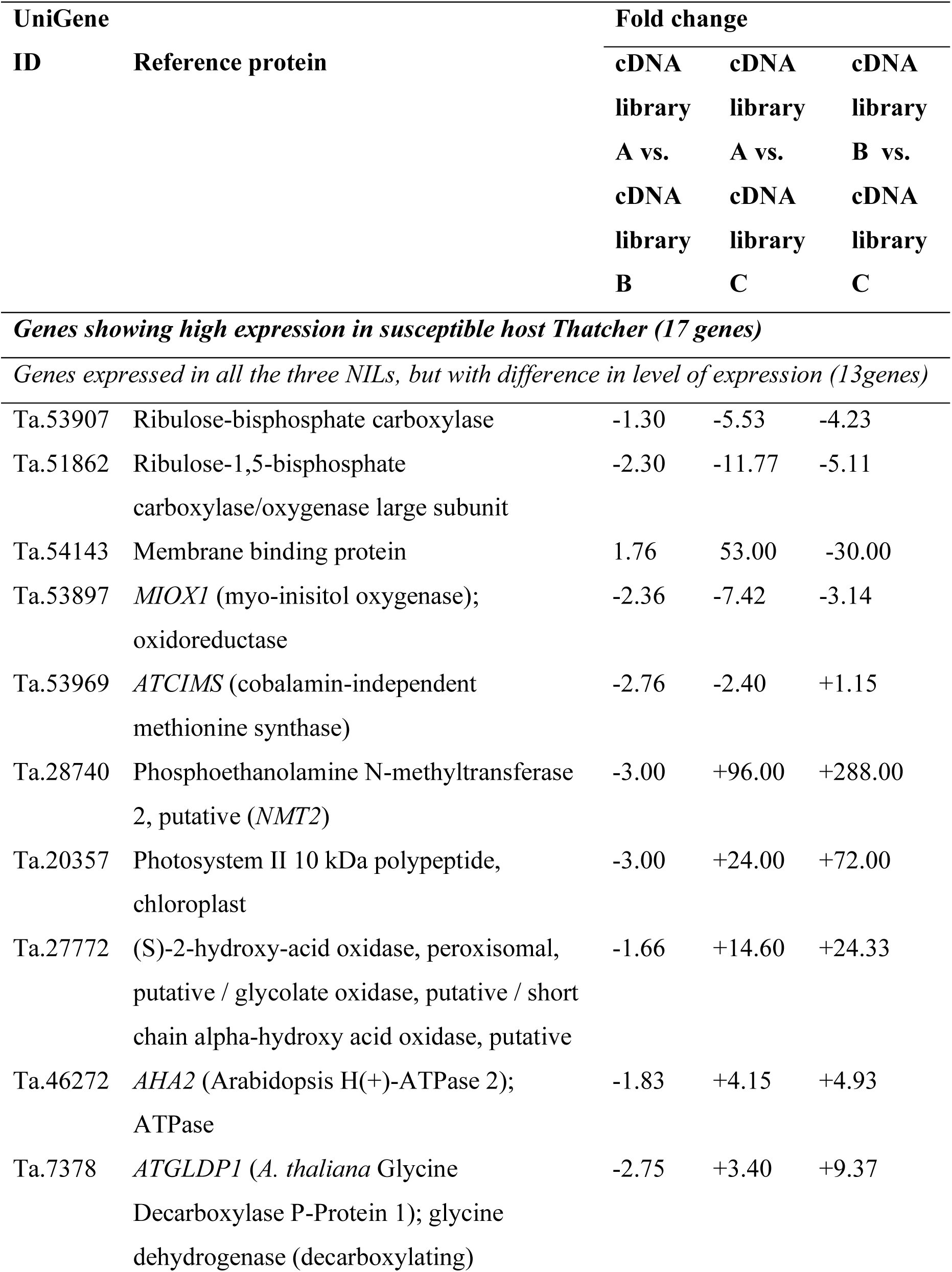
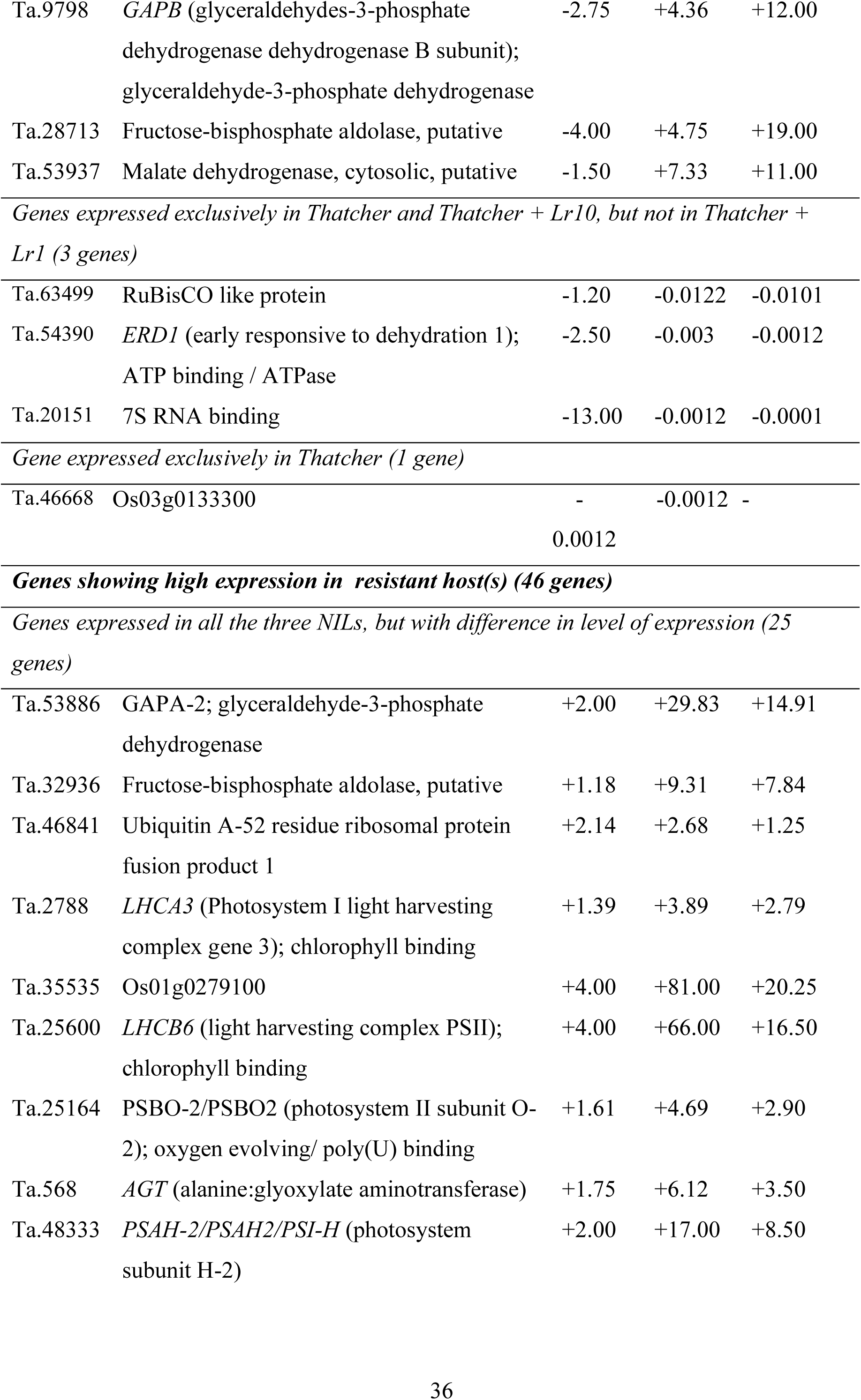
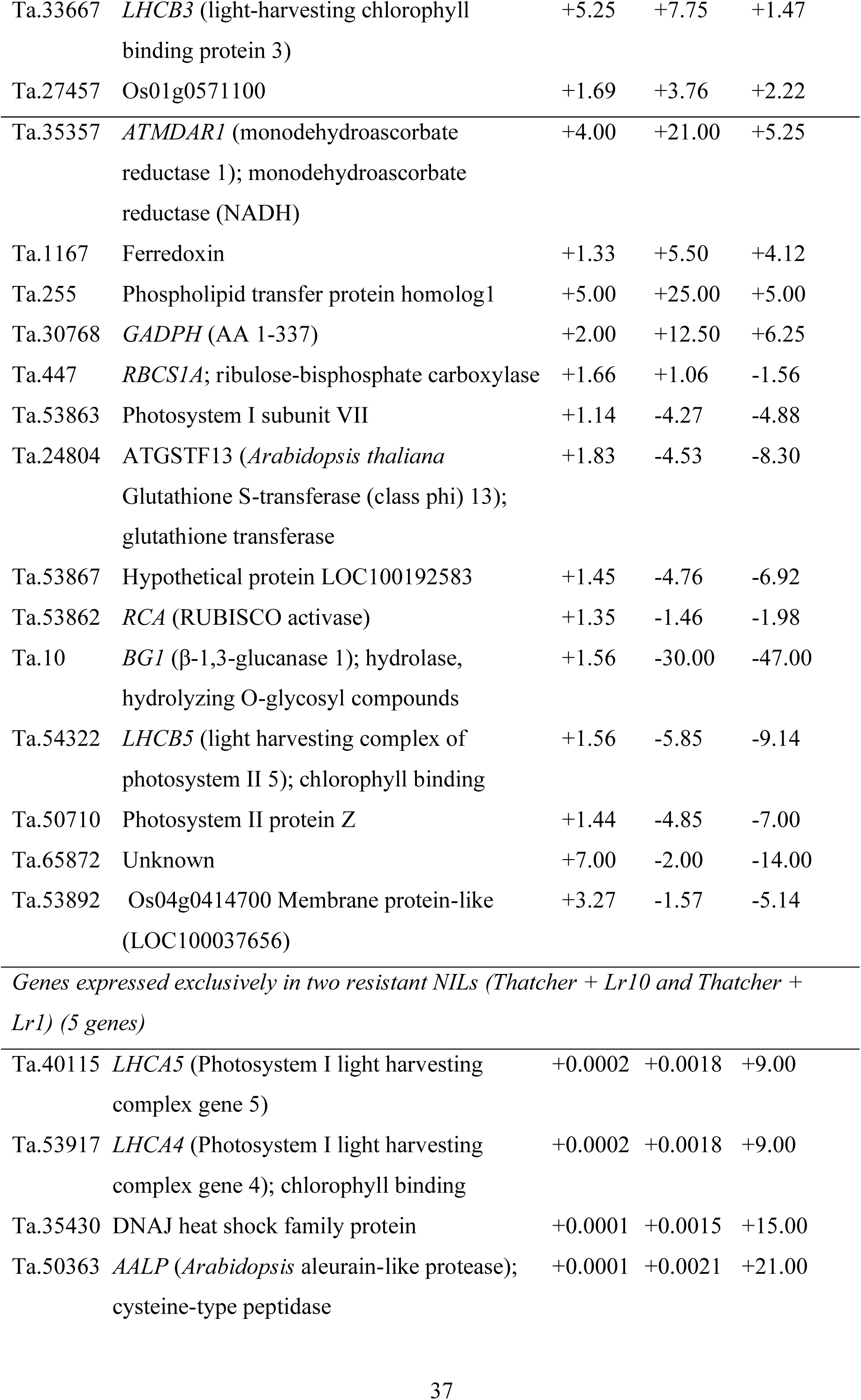
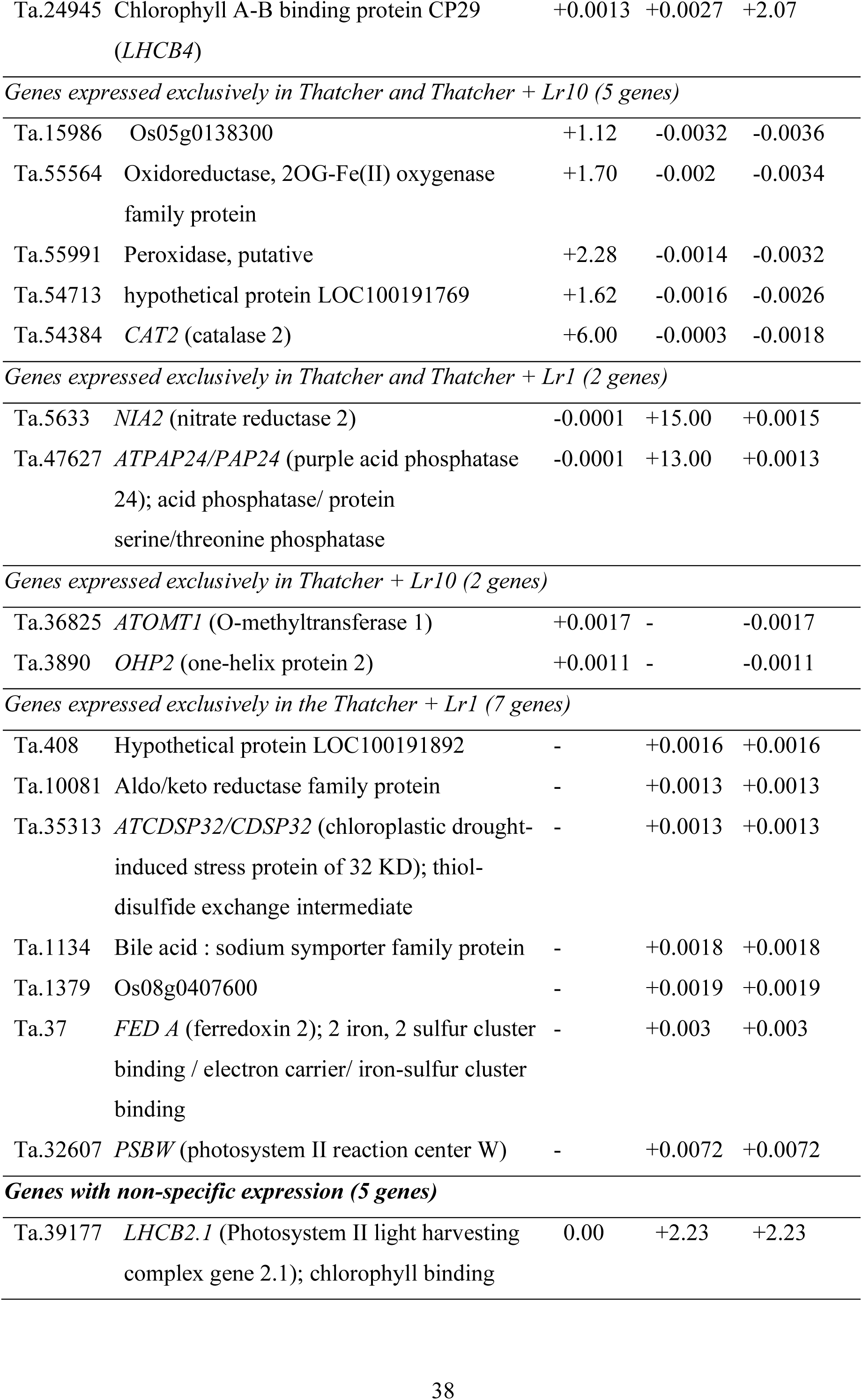
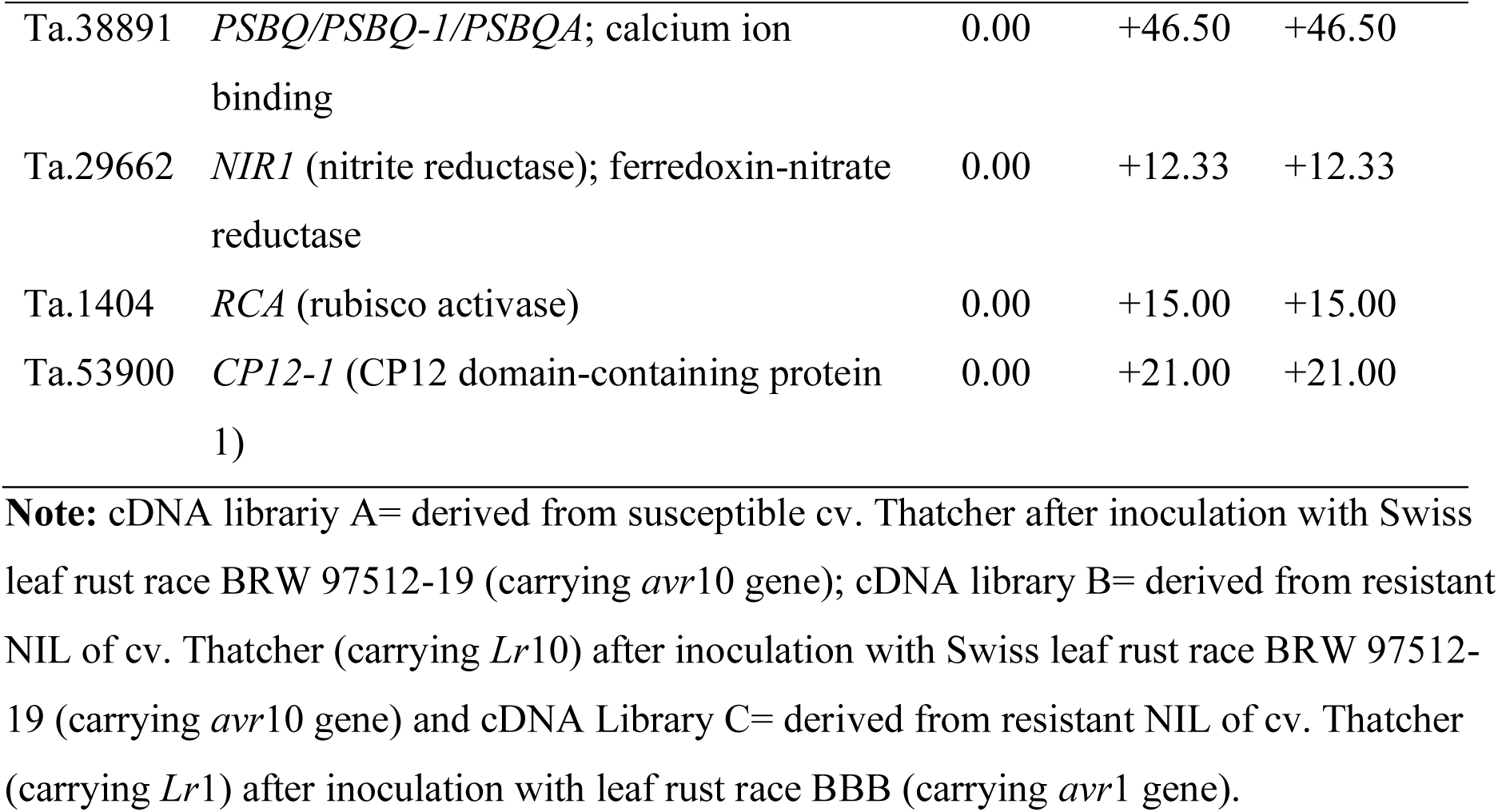
Details of wheat UniGenes appeared in different libraries and fold change in their abundance.

### Genes showing high expression in susceptible host Thatcher

*Genes expressed in all three NILs, but with differences in the level of expression*: In the susceptible host Thatcher, 1UniGenes were found to exhibit high expression. Among these UniGenes, some encode proteins such as Ribulose-1,5-bisphosphate carboxylase/oxygenase (*RuBisCO*) large subunit, membrane binding protein, *MIOX1* (myo-inisitol oxygenase1) which is involved in cell wall metabolism, and *ATCIMS* (cobalamin-independent methionine synthase) which is involved in methionine metabolism, as shown in Table 4.

The UniGene encoding caffeic acid 3-O-methyltransferase (*COMT*) is known to play an important role in lignification at the infection site in tobacco plants (Guo et al. 2001). This enzyme is involved in the process of callose deposition in plasmodesmata, which modifies cell wall architecture through crosslinking and lignification events. As a result, both symplastic and apoplastic routes for signal molecules become blocked, leading to the progressive shutdown of defense signaling pathways (Hammond-Kosack and Jones, 1996). Therefore, the gene encoding *COMT* is essential for regulating the host response following rust infection.

Previous studies have shown that the breakdown of photosynthetic components, such as the large subunit of *RuBisCO*, by the enzymes belonging to the infecting pathogen can result in sub-optimal functioning of photosynthesis in susceptible plants (Van der Westhuizen and Botha 1993). In the present study, we observed abundant expression of the gene encoding the large subunit of *RuBisCO* in susceptible plants, possibly due to the induction of PAMP-triggered immunity (PTI). PTI results from the early identification of pathogen-associated molecular patterns (PAMPs) by the pattern-recognition receptors (PRRs) of host cells (Pieterse et al. 2009). It is plausible that PRRs shield the photosynthesis machinery of the host cell from degradation by the enzymes belonging to the infecting pathogen.

The results of our study suggest that changes in gene expression in response to leaf rust infection are widespread and involve a variety of biological processes. Besides the genes mentioned above, several other UniGenes with high expression in susceptible interaction encode important proteins, including membrane binding protein, Phosphoethanolamine N-methyltransferase 2 (*NMT2*), and *AHA2* (*Arabidopsis* H(+)-ATPase 2) (Table 4). These proteins belong to pathways involved in disease/defense, photosynthesis, glycolysis, and oxidation, indicating alterations in different biological processes during compatible interactions.

The expression of photosynthetic-related genes in wheat, such as *rbcl*, may be influenced by various factors, including the accumulation of leaf soluble sugar (Cheng et al. 1998). Lacoc et al. (2003) also observed an abundance of *RuBisCo* in aphid-infected wheat plants, possibly indicating the role of pattern-recognition receptors (PRRs) in protecting the photosynthesis machinery of the host cell from degradation by the pathogen’s enzymes. *MIOX1* (myo-inositol oxygenase 1), belonging to the myo-inositol oxygenase gene family, has been reported to be involved in changing the redox status and polysaccharides of the cell wall in leaf rust-infected host cells (Kanter et al. 2005). Additionally, the gene encoding *ATCIMS* (cobalamin-independent methionine synthase) may be a new candidate for regulating the resistance and susceptibility in wheat plants.

Recent studies have highlighted the role of certain genes in regulating resistance to fungal infection. For example, the *Arabidopsis* gene *AtPEN1* has been reported to provide non-host resistance to the fungus *Blumeria graminis* f. sp hordei (Hua et al. 2009). *AtPEN1* encodes a SNARE receptor family protein AtSYP121, which mediates the formation of papillae in the pathogen invasion site by regulating vesicle traffic and the signal pathway. The Atpen1-1 mutant lacking non-host resistance showed S-nitrosylational modification of their protein product *AtSYP121* by cobalamin-independent methionine synthase (*CIMS*), indicating its role in post-translational modification of resistance gene products and regulating gene function.

In addition to these genes, other UniGenes that showed high expression in susceptible interactions encoded important proteins like membrane binding protein, Phosphoethanolamine N-methyltransferase 2 (*NMT2*), and *AHA2* (*Arabidopsis* H(+)-ATPase 2) (Table 4). These proteins largely belong to disease/defense, photosynthesis, glycolysis, and oxidation pathways, indicating alterations of different biological processes during compatible interactions. Therefore, understanding the expression and function of these genes may provide insights into the mechanisms underlying host-pathogen interactions in wheat.

*Genes expressed exclusively in Thatcher and Thatcher + Lr10, but not in Thatcher + Lr1:* The UniGenes identified in this study encode for several important proteins, including RuBisCO-like protein, *ERD1*, and a protein similar to 7S RNA binding domain (Table 4). Following leaf rust infection in susceptible wheat plants, the expression of *ERD1* and the gene encoding RuBisCO-like protein was observed to be high, indicating that they were induced as a consequence of dehydration stress aroused during a compatible interaction. *ERD1* encodes for the Clpa protein, which is involved in ATP binding/ATPase activity and plays a role in stress response (Hong et al. 2003). Additionally, disease resistance protein SRP54, which contains a 7S RNA binding domain involved in SRP-dependent translational protein targeting to the membrane, was found to be highly expressed in susceptible interaction. *SRP54* is an important component of the signaling pathway and its high expression in susceptible interaction suggests modulation of a disease resistance pathway by an unknown mechanism. All these proteins belong to defense, energy production, and protein synthesis machinery, highlighting the alterations of different biological processes during compatible interactions.

*Gene expressed exclusively in Thatcher*: It is interesting to note that the expression of UniGene Ta.46668, which encodes a hypothetical protein of *O. sativa*, was detected exclusively in Thatcher and not in Thatcher + Lr10 or Thatcher + Lr1. This suggests that this gene may play a role in susceptibility or be completely repressed in resistant lines after inoculation. Therefore, it may be a new candidate gene responsible for the compatible susceptible response.

The high expression of these UniGenes in the susceptible host Thatcher suggests that the molecular changes initiated in susceptible reactions involve various biological processes such as energy production, photosynthesis, membrane binding, glycolysis, and oxidation. However, the precise functions of these genes in susceptibility to leaf rust in wheat are still not completely understood and further research is needed to fully elucidate their roles in the susceptible response.

### Genes showing high expression in resistant host(s)

*Genes expressed in all three NILs, but with a difference in the level of expression*: It may be recalled that 25 UniGenes (other than those 13 UniGenes, which expressed in all three cDNA NILs with high expression in susceptible NIL or library A) had high expression in Thatcher + *Lr*10 or in Thatcher + *Lr*1 or in both these resistant NILs. These UniGenes encode important proteins like glutathione S-transferase (class phi), RuBisCO activase, β-1,3-glucanase 1, and many other proteins (Table 4). Glutathione S-transferase (*GST*) belongs to the family of enzymes that play a key role in detoxifying xenobiotic compounds in plants. The high expression of the *GST* gene in resistant NILs Thatcher + Lr10 and Thatcher + Lr1 indicates its involvement in the defense response against leaf rust. Previous studies have shown that *GST* expression is induced upon pathogen attack, which is consistent with the activation of the plant defense response (Roxas et al., 1997). In addition, the production of salicylic acid has been shown to correlate with *GST* expression (Dixon et al., 2010). Dixon et al. (2010) also highlight the important roles that glutathione transferases play in plant secondary metabolism. Therefore, it is suggested that *GST*s have a vital role in plants’ response to pathogen attack and their defense mechanisms. RuBisCO activase, on the other hand, is an enzyme that plays a crucial role in the process of photosynthesis by catalyzing the activation of RuBisCO. The high expression of RuBisCO activase in leaf rust-inoculated wheat plants can be attributed to the increased activation of RuBisCO, which helps in maintaining the photosynthetic potential of infected leaves. This is an important defense mechanism in plants, as photosynthesis provides the energy necessary for the plant to mount an effective defense response. β-1,3-glucanase 1 is a protein that regulates plant symplasmic permeability via hydrolysis of callose in plasmodesmata. It plays a role in both development and defense processes in plants. The high expression of this gene in resistant NILs Thatcher + Lr10 and Thatcher + Lr1 suggests its involvement in the defense response against leaf rust. This is supported by previous studies that have shown that β-1,3-glucanase expression is induced upon pathogen attack and is involved in the degradation of fungal cell walls, which helps in limiting the spread of the pathogen. In our study, we analyzed the transcriptome of three near-isogenic lines (NILs) with different combinations of Lr10 and Lr1 genes in response to *Puccinia triticina*, the causal agent of wheat leaf rust disease. Our analysis revealed that a group of UniGenes exhibited expression in all three NILs, with a particularly high expression in the resistant NILs. These UniGenes were found to be involved in the regulation of photosynthesis machinery, ion fluxes, and reactive oxygen species (ROS) production. Furthermore, we found that the UniGenes identified in our study appeared to be specific to resistance reactions involving Lr10 and/or Lr1 genes. Our findings indicate that the expression of certain UniGenes was differentially regulated by the two different Lr genes, namely Lr10 and Lr1 in the common genetic background. Interestingly, some of these UniGenes exhibited low expression in Thatcher + Lr1, suggesting that the presence of Lr1 may modulate the expression of these genes.

*Genes expressed exclusively in Thatcher + Lr10 and Thatcher + Lr1*: Our study identified five UniGenes that were expressed in both resistant NILs (Thatcher + Lr10 and Thatcher + Lr1) but not in the susceptible NIL Thatcher (Table 4). One of these UniGenes, Ta.50363, encodes Triticain gamma (CTSH), a protein similar to Cathepsin H, a lysosomal cystein protease involved in intracellular degradation. CathB-like proteases have been shown to trigger a hypersensitive response in plants by co-expression with specific effectors and their cognate R genes, suggesting that Ta.50363 may play a role in the hypersensitive response to leaf rust disease. Another UniGene identified in our study, Ta.35430, encodes a protein similar to heat shock family protein DnaJ, which interacts with the chaperone HSP70-like DnaK protein in prokaryotes. In eukaryotes, HSP90 and HSP70 are important components of a multi-chaperone system that co-operates with co-chaperones like HSP40 (a DnaJ homolog). Thus, Ta.35430 may be involved in conferring tolerance to abiotic stresses in wheat. The remaining three UniGenes, Ta.24945, Ta.40115, and Ta.53917, encode proteins similar to light-harvesting complex proteins of *A. thaliana*. These UniGenes may play a role in photosynthesis machinery, a key process in plant metabolism, and their specificity to leaf rust resistance makes them particularly interesting. Overall, our study highlights the importance of these five UniGenes in conferring resistance to leaf rust disease in wheat.

*Genes expressed exclusively in Thatcher and Thatcher + Lr10*: Five UniGenes were found to be expressed exclusively in Thatcher and Thatcher + Lr10, with a higher expression level in the resistant NIL (Table 4). Of these, Ta.55564 encodes a 2OG-Fe(II) oxygenase, which is involved in iron ion and L-ascorbic acid binding, as well as oxidoreductase activity (Damme et al. 2008). *AlkB*, a member of the 2OG-Fe(II) oxygenase superfamily, has been shown to catalyze the oxidative detoxification of alkylated bases, indicating its potential role in disease resistance by altering the redox status of infected cells. Another highly expressed UniGene, Ta.54384, encodes catalase 2, which plays a crucial role in systemic acquired resistance in *A. thaliana* (Frugoli et al. 1996). Catalases are important in H2O2 scavenging in different plant cell compartments and are involved in defense gene activation in response to the production of active oxygen species, such as H2O2, in disease-infected cells (Levine et al. 1994). UniGene Ta.55991, which encodes peroxidases, also showed high expression in the resistant NIL Thatcher + Lr10. Peroxidases are part of the pathogenesis-related (PR) protein family and play a role in restricting fungal development through hydrolytic action on their cell walls (Klessig and Malamy 1994, Hunt and Ryals 1996, Kombrink and Somssich 1997, Nandi et al. 2004). The altered redox status of infected cells of resistant hosts might activate the peroxidase cycle-based hypersensitive response (Singh et al. 2012). The interaction between SA and catalase or ascorbate peroxidase in infected resistant cells results in the formation of phenolic free radicals, which are proposed to be involved in the induction of systemic acquired resistance (SAR) (Durner and Klessig 1995). Thus, the high expression of UniGenes encoding catalase 2 and peroxidase signifies their potential role in the induction of SAR. The functions of the remaining two highly expressed UniGenes remain unknown. The absence of the expression of these five UniGenes in Thatcher + Lr1 is not surprising, as their expression seems to depend on the interaction between Thatcher NILs (both susceptible and resistant (Lr10)) and the leaf rust race BRW 97512-19. These genes were not specifically associated with Lr1 gene expression and were not induced by the leaf rust race BBB.

*Genes expressed exclusively in Thatcher and Thatcher + Lr1:* Two UniGenes, Ta.5633 and Ta.47627, exhibited exclusive expression in the Thatcher and Thatcher + Lr1 cultivars, with relatively high expression levels observed in Thatcher + Lr1. The UniGene Ta.5633 exhibited significant similarity to the *Nia2* gene, which encodes nitrate reductase (NR) (Table 4). NR is a key enzyme involved in the synthesis of nitric oxide (*NO*) during host-pathogen interactions (Oliveira et al., 2010). *NO* regulates the expression of resistance gene products via S- nitrosylation, which is critical for plant defense against pathogens. The second UniGene, Ta.47627, encodes the purple acid phosphatase 24 (*PHP24*) protein (Table 4). Recent studies have shed more light on the structure and catalysis of PHPs, emphasizing their significance in various cellular processes. For example, Chiara et al. (2019) provide an overview of the structure and catalysis of alkaline phosphatase, highlighting its diverse applications. Additionally, König et al. (2020) discovered that the alkaline phosphatase Pho91 acts as a multicopy suppressor of the plasma membrane proton ATPase Pma1 in Saccharomyces cerevisiae, indicating its crucial role in maintaining cellular homeostasis. These recent studies further highlight the importance of PHPs in various cellular processes and their potential applications.

*Genes expressed exclusively in Thatcher + Lr10*: Our analysis revealed the expression of two UniGenes, Ta.36825 and Ta.3890, in the Thatcher + Lr10 cultivar. Ta.36825 showed significant similarity to the *ATOMT1* (O-methyltransferase1) gene of *Arabidopsis thaliana*, which is involved in several important biological processes such as lignin formation, biosynthesis of sinapate esters, and cell wall digestibility (Goujon et al., 2003). The expression of Ta.36825 suggests its potential role in similar processes in wheat. The second UniGene, Ta.3890, encodes a protein similar to *OHP2* (one-helix protein 2) of *A. thaliana*, which is known to be triggered by light stress, and its transcripts and proteins accumulate in a light intensity-dependent manner (Andersson et al., 2003). The expression of Ta.3890 in Thatcher + Lr10 suggests a potential involvement in the response to light stress. Interestingly, the expression of both UniGenes appears to be associated with the leaf rust resistance gene Lr10. This finding suggests a possible role of Ta.36825 and Ta.3890 in the defense mechanisms against the leaf rust pathogen in Thatcher + Lr10.

*Genes expressed exclusively in the Thatcher + Lr1*: Seven UniGenes that are exclusively expressed in Thatcher + Lr1 (Table 4), encoding proteins that are similar to five known proteins of *A. thaliana* and two unknown proteins of *O. sativa*. Among these UniGenes, Ta.32607 encodes the PSBW protein, which is a crucial component of PS-II and plays a vital role in the regulation and biogenesis of the photosynthetic apparatus. Plants lacking PSBW proteins have been observed to exhibit a reduction of approximately 40% in PSII and the oxygen-evolving complex, as reported in Swiss-Prot. Moreover, the UniGene Ta.37 encodes the FED A protein that participates in a range of redox reactions in plants (Somers et al. 1990), and UniGene Ta.35313 encodes a protein that is similar to the drought-induced proteins.

The observed differential gene expression after pathogen inoculation suggests that genes involved in photosynthesis, metabolism, and energy production are reprogrammed during the resistance reaction to fulfill the energy demand created during the hypersensitive reaction. This is consistent with previous studies that showed that resistance reactions involve a cascade of protein phosphorylation, ion fluxes, and reactive oxygen species (ROS) (Manickavelu et al. 2010). In another study on Lr34-mediated leaf rust resistance in wheat, Bolton et al. (2008) reported that resistance response imposes a high energy demand that leads to the induction of multiple metabolic responses to support cellular energy requirements. They also suggested that a wide variety of genes were reprogrammed during resistance reactions. Indeed, high expression of various housekeeping genes (besides various defense-related genes) is likely to accompany the host defense response to ensure that ample pools of precursor compounds are maintained (Kawalleck et al. 1992).

In addition, co-regulation of genes involved in different stress-related signaling pathways, such as DnaJ, suggests cross-talk between pathways and results in cross-tolerance of the host genotype (Manickavelu et al. 2010). Cross-tolerance is a biological phenomenon where a host develops resistance to one type of stress if it is already tolerant to another form of stress (Tuteja et al. 2007). In tomato plants, Capiati et al. (2006) found that wounding can increase salinity tolerance as a result of cross-tolerance.

### Genes with non-specific expression

Five (5) UniGenes (Ta.39177, Ta.38891, Ta.29662, Ta.1404, and Ta.53900) showed the same level of expression in Thatcher and Thatcher + Lr10 and a slightly high expression in Thatcher + Lr1 (Table 4) and are related to functions such as photosynthesis, calcium ion binding, nitrate pathway, oxidation, and Calvin cycle regulation. However, it appears that their high expression is not inducible by the interaction between Thatcher + Lr1 and leaf rust race BBB, and no role in resistance induction was observed.

### Clustering and co-expression of differentially expressing UniGenes in different libraries

During resistance, a cell at the site of infection initiates a cascade of rapid biological functions and processes that are controlled by different co-regulated genes. This leads to the sequential expression of genes involved in different biochemical pathways, which collectively constitute a signal transduction pathway (Jiang et al. 2007).

In the present study, 68 differentially expressed UniGenes were clustered into five groups. The UniGene database provided functional annotation for several of these genes (Table 4). The identified five main clusters (Clusters I, II, III, IV, and V), with each cluster representing a group of UniGenes that expressed similarly (Table 5). The number of UniGenes present in each cluster varied from 3 (cluster I) to 26 (cluster V).

**Table 5.** Grouping of 68 differentially expressing UniGenes of wheat in five clusters (I-V)

| UniGenes | Cluster |
| --- | --- |
| Ta.53969 | I |
| Ta.53817, Ta.40115, Ta.54390, Ta.24945, Ta.20151, Ta.50363, Ta.63499, Ta.5633, Ta.255, Ta.35357, Ta.27457, Ta.46668, Ta.25600, Ta.568, Ta.25164, Ta.35535, Ta.33667, Ta.10061, Ta.46841 | II |
| Ta.65872, Ta.55992, Ta.53862, Ta.54713, Ta.55991, Ta.54384, Ta.55564, Ta.15986, Ta.447, Ta.3490 | III |
| Ta.24804, Ta.50710, Ta.54322, Ta.53367, Ta.10, Ta.37, Ta.53863, Ta.53897, Ta.1379, Ta.51862, Ta.54163, Ta.53907 | IV |
| Ta.47627, Ta.35430, Ta.1167, Ta.1134, Ta.53886, Ta.30768, Ta.48333, Ta.2788, Ta.406, Ta.32936, Ta.1404, Ta.29662, Ta.53900, Ta.38891, Ta.39177, Ta.28713, Ta.7378, Ta.9798, Ta.36825, Ta.20357, Ta.53937, Ta.35313, Ta.27772, Ta.32607, Ta.28740, Ta.46272 | V |

*Genes within a cluster have similar expression patterns and diverse functions*: The UniGenes grouped together in each cluster exhibited diverse roles in cellular processes. Cluster I included genes involved in nitrate reduction and phospholipid transfer protein, while cluster II included genes encoding ubiquitin A-52 residue ribosomal protein and other proteins. Cluster III comprised genes related to glycolysis, tricarboxylic acid cycle, oxidative phosphorylation, and heat shock proteins. Cluster IV was dominated by photosynthesis-related genes, as well as genes involved in respiration, ion fluxes, and stress response. Finally, cluster V contained genes encoding proteins such as aldo/keto reductase family protein, RuBisCo activase, peroxidase, catalase, oxidoreductase, chloroplastic drought-induced stress protein, and ribulose biphosphate carboxylase. (Table 5)

The functional annotation of all clustered UniGenes is somewhat limited, particularly in the case of wheat, making it challenging to assign a function to each of the clustered genes. Nonetheless, it is evident that the genes within a cluster had diverse functions, and genes with similar functions were often found in different clusters. Thus, co-expression of differentially expressing UniGenes could potentially lead to a coordinated defense response of the host plant against the pathogen. In previous studies, genes belonging to the above functional categories also demonstrated differential expression following leaf rust inoculation of resistant wheat cultivars (Fofana et al. 2007, Hulbert et al. 2007, Manickavelu et al. 2010, Dhariwal et al. 2011, Singh et al. 2012).

### Clustering and co-expression of differentially expressing UniGenes in different tissues

Based on the analysis of the differentially expressed UniGenes, it was observed that all of them had high expression levels in leaves, with moderate expression in sheath, stem, and inflorescence, and low expression in seed, root, flower, crown, callus, and cell culture. Although none of the genes were exclusively expressed in leaves, one UniGene (Ta.53892) was found to be absent in leaves but was expressed in leaf sheath and stem, as well as other tissues. It is worth noting that wheat leaves are the primary target tissue for leaf rust infection and fungal growth, as well as for resistance response such as programmed cell death at the infection site. The expression pattern of the identified UniGenes suggests that they are directly or indirectly involved in the hypersensitive resistance response following fungal infection. The moderate to low expression of these genes in other tissues suggests that the reprogramming of targeted cells also leads to a secondary response, known as systemic acquired resistance, in other vegetative tissues (Dempsey et al. 1994).

Based on the findings of our study and information from previously published literature (Liebenberg 2007, Fofana et al. 2007, Hulbert et al. 2007, Manickavelu et al. 2010, Dhariwal et al. 2011, Singh et al. 2012), we developed a simple model that illustrates the mechanisms of leaf rust resistance and susceptibility in wheat. Our model demonstrates how host cells respond and coordinate the actions of numerous genes that are activated after pathogen recognition by Lr genes (as shown in Fig. 1), leading to either resistance or susceptibility.

**Fig. 1.**
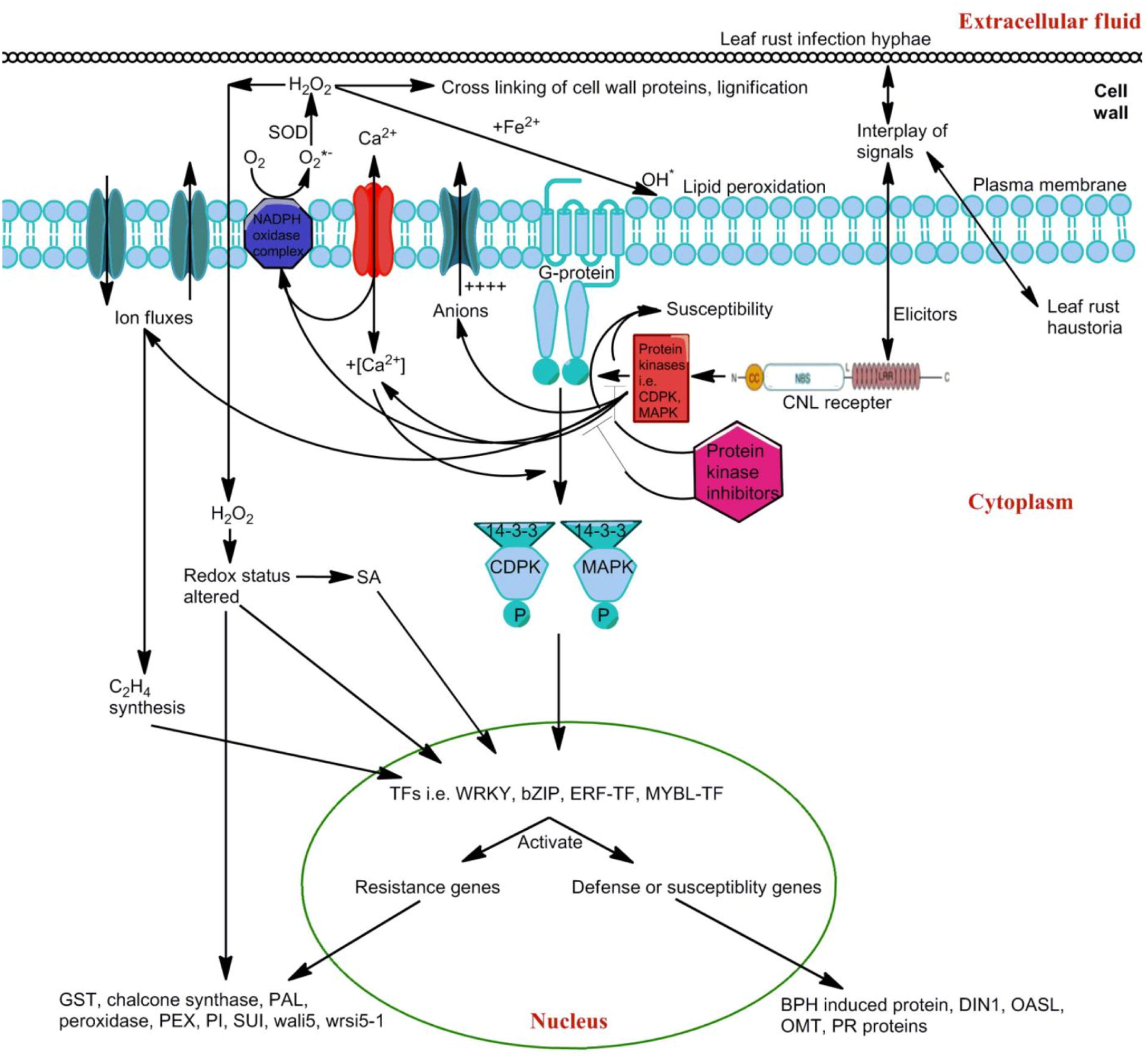
A model of host resistance and susceptibility responses in leaf rust infected wheat plants. Resistance response was mediated by seedling resistance gene (e.g. *Lrl* or *LrlO)* which encodes CC-NBS-LRR domain protein. The genes identified in the present study were involved in various processes like Ca^2^+ ion binding, ion fluxes, alteration of redox status, lignifications, target delivery of pathogenesis related proteins etc. *SOD:* Superoxidase; *CDPK:* cyclin­ dependent protein kinases, MAPK: mitogen-activated protein kinase, *CNL* receptor: coiled coil­ nucleolar binding site-leucine-rich repeats receptor, SA: salicylic acid, *GST:* glutathione S - transferase; PAL: phenylalanine arnmonia-lyase, *PEX:* peroxidase, *P*/, proteinase inhibitors, *BPH* induced proteins: brown plant hopper induced proteins, *DINI:* dark-inducible protein, *OASL:* O-acetylserine(thiol)lyase, *OMT:* O-methyltransferase.

### Gene expression pattern validation

Twenty-one UniGenes out of the 68 DEUs were matched with individual wheat probes on the 61K platform in PLEXdb. Among these, 14 (∼67%) of the 21 matched UniGenes showed similar expression patterns in both the DDD analysis and microarray analysis (as indicated in Table 6). In the DDD analysis, 13 of the 21 UniGenes exhibited maximum expression in the resistant cultivars (Thatcher + Lr10 and Thatcher + Lr1), and 8 (∼62%) of these 13 UniGenes also displayed maximum expression in the resistant cultivar using microarray experimental data. Moreover, approximately 80% and ∼67% of UniGenes belonging to the ‘susceptible’ and ‘non-specific’ categories, respectively, also demonstrated similar expression patterns using DDD analysis and microarray experimental data. Interestingly, out of the 21 UniGenes mentioned above, 9 (∼43%) UniGenes showed the same tissue-specific expression pattern using both DDD analysis and microarray experimental data, highlighting the significance of PLEXdb in validating the gene expression results.

**Table 6.** Details of comparison of expression pattern of differentially expressed wheat UniGenes between DDD analysis and microarray experiment.

| UniGene ID | Maximum expression in resistant vs. susceptible NILs |  |  | Maximum expression in tissue |  |
| --- | --- | --- | --- | --- | --- |
|  | Using DDD | Using microarray |  | Using DDD | Using microarray |
| Ta.53863 | Resistant | Resistant |  | Stem | Seedling: leaf |
| Ta.28740 | Susceptible | Resistant |  | Stem | Seedling: leaf |
| Ta.63499 | Susceptible | Susceptible |  | Leaf | Seedling: leaf |
| Ta.51862 | Susceptible | Susceptible |  | Leaf | Seedling: leaf |
| Ta.38891 | Equal | Equal |  | Leaf | Seedling: leaf |
| Ta.35535 | Resistant | Resistant |  | Leaf | Seedling: leaf |
| Ta.53867 | Resistant | Susceptible |  | Stem | Seedling: leaf |
| Ta.46272 | Resistant | Resistant |  | Leaf | Floral bracts: before anthesis |
| Ta.53907 | Susceptible | Susceptible |  | Sheath | Seedling: leaf |
| Ta.53862 | Resistant | Susceptible |  | Stem | Seedling: leaf |
| Ta.25164 | Resistant | Resistant |  | Leaf | Seedling: leaf |
| Ta.10 | Resistant | Resistant |  | Leaf | Floral bracts: before anthesis |
| Ta.29662 | Equal | Resistant |  | Leaf | Seedling: leaf |
| Ta.1404 | Equal | Equal |  | Leaf | Seedling: leaf |
| Ta.50710 | Resistant | Susceptible |  | Leaf | 22 DAP endosperm |
| Ta.50363 | Resistant | Susceptible |  | Leaf | Seedling: leaf |
| Ta.1167 | Resistant | Resistant |  | Leaf | Seedling: leaf |
| Ta.53892 | Resistant | Resistant |  | Sheath | Seedling: leaf |
| Ta.255 | Resistant | Resistant |  | Stem | Seedling: leaf |
| Ta.30768 | Resistant | Susceptible |  | Cell culture | 22 DAP endosperm |
| Ta.53937 | Susceptible | Susceptible |  | Crown | 22 DAP endosperm |

### In-silico physical mapping of validated UniGenes on wheat chromosomes

The mapping process involved an in-silico approach using a BLAST search against previously mapped ESTs/EST contigs (as shown in Table 7). Out of the 21 UniGenes that matched with wheat microarray probes, 11 (∼52%) UniGenes were assigned to 26 loci on 13 individual wheat chromosomes (as indicated in Table 7). The distribution of loci on the seven homoeologous groups and the three subgenomes was found to be nonrandom (P < 0.02). Both chromosomal arms had an equal number of loci, with more than one-third of the loci mapped to the centromeric bins of chromosomes, which was disproportionate relative to the differences in DNA content and arm length (as shown in Table 7). It is worth noting that due to the very low number of loci in the present study, these results are not consistent with an earlier study (Mohan et al. 2007) which reported a higher number of loci being mapped on long arms, with more loci mapped to the distal regions of chromosome arms. A fairly large proportion (∼81%; 9 out of 11) of UniGenes had ≥2 loci, which were mostly (∼66%; 6 out of 9) homoeoloci (as indicated in Table 7). Interestingly, one copy (locus) of each of the two UniGenes, namely Ta.50363 and Ta.53863, were mapped to chromosome 5DL, which is the carrier chromosome arm of the leaf rust resistance gene Lr1. These physical mapping results could be useful for comparative genomics, gene cloning, and marker-assisted selection in resistance breeding programs.

**Table 7.** Details of wheat UniGenes assigned to deletions bin(s) in wheat chromosome(s).

| **UniGene ID** | **Chromosomal position** | **Position in bin(s)on respective chromosome(s)** |
| --- | --- | --- |
| Ta.53863 | 5AL, 5BL, 5DL | C-5AL10-0.57, C-5BL14-0.75, C-5DL1-0.60 |
| Ta.28740 | 1BL, 1DL, 4BS | 1BL1-0.47-0.69, 1DL2-0.41-1.00, 4BS1-0.81-1.00 |
| Ta.53867 | 6BL, 6DL | 6BL3-0.36-0.40, C-6DL6-0.29 |
| Ta.46272 | 5AS, 5BS, 5DS | 5AS7/10-0.98-1.00, 5BS6-0.81-1.00, 5DS2-0.78-1.00 |
| Ta.53862 | 4DS | C-4DS1-0.53 |
| Ta.10 | 1BL, 1DL, 7BS | C-1BL6-0.32, 1DL2-0.41-1.00, C-7BS1-0.27 |
| Ta.1404 | 4DS | C-4DS1-0.53 |
| Ta.50363 | 5AL, 5DL | 5AL10-0.57-0.78, 5DL1-0.60-0.74 |
| Ta.1167 | 6AS, 6BS, 6DS | 6AS5-0.65-1.00, 6BS-Sat, 6DS6-0.99-1.00 |
| Ta.255 | 3BS, 3DS | 3BS8-0.78-1.00, 3DS6-0.55-1.00 |
| Ta.30768 | 4DS, 6AL, 6BL | 4DS3-0.67-0.82, C-6AL4-0.55, C-6BL3-0.36 |

## Conclusions

This study utilized NCBI’s Digital Differential Display tool and Cluster 3.0 program to compare the transcriptome of leaf rust-infected susceptible and resistant wheat NILs, providing novel insights into the genes involved in resistance response during leaf rust infection. The identification of known and unknown genes associated with resistance response highlights the potential of these tools in transcriptomic analysis. The co-regulation of genes with diverse functions during infection suggests a coordinated response by the host plant. The high expression of differentially expressed genes in leaves and validation using microarray gene expression data further support the involvement of these genes in the defense response. The mapping of validated UniGenes to wheat chromosomes enhances our understanding of the wheat genome. This study has implications for future genome-wide transcriptomic analyses in response to biotic and abiotic stresses and could aid in the development of molecular markers and genome editing for the creation of leaf rust-resistant wheat cultivars.

## Acknowledgement

ChatGPT was used to improve grammar of this manuscript.

